# Detection of a novel insect specific flavivirus across ecologically diverse populations of *Aedes aegypti* on the Caribbean Island of Saint Lucia

**DOI:** 10.1101/434183

**Authors:** C.L. Jeffries, M. White, L. Wilson, L. Yakob, T. Walker

**Affiliations:** Department of Disease Control, London School of Hygiene and Tropical Medicine, London, WC1E 7HT, United Kingdom

## Abstract

Outbreaks of mosquito-borne arboviral diseases including dengue virus (DENV), Zika virus (ZIKV), yellow fever virus (YFV) and chikungunya virus (CHIKV) have recently occurred in the Caribbean. The geographical range of the principle vectors responsible for transmission, *Aedes (Ae.) aegypti* and *Ae. albopictus* is increasing and greater mosquito surveillance is needed in the Caribbean given international tourism is so prominent. The island of Saint Lucia has seen outbreaks of DENV and CHIKV in the past five years but vector surveillance has been limited with the last studies dating back to the late 1970s. Natural disasters have changed the landscape of Saint Lucia and the island has gone through significant urbanisation. In this study, we conducted an entomological survey of *Ae. aegypti and Ae. albopictus* distribution across the island and analysed environmental parameters associated with the presence of these species. Although we collected *Ae. aegypti* across a range of sites across the island, no *Ae. albopictus* were collected despite traps being placed in diverse ecological settings. The number of *Ae. aegypti* collected was significantly associated with higher elevation and semi-urban settings yielded female mosquito counts per trap-day that were 5-fold lower than urban settings. Screening for arboviruses revealed a high prevalence of a novel insect-specific flavivirus closely related to cell fusing agent virus (CFAV). We discuss the implications that natural disasters, water storage and lack of mosquito surveillance have on arboviral outbreaks in Saint Lucia and implications for insect only flaviviruses on surveillance and detection of pathogenic flaviviruses.

## INTRODUCTION

Medically important arboviruses that cause human morbidity and mortality are predominantly transmitted by mosquitoes. There are more than 600 known arboviruses and related zoonotic viruses with more than 80 known to be human pathogens. Outbreaks of dengue virus (DENV), Zika virus (ZIKV), yellow fever virus (YFV) and chikungunya virus (CHIKV) are increasing ^1^ and there is potential for zoonotic viruses to spill-over into human populations. Arboviral disease transmission mostly occurs in tropical countries of Southeast Asia and South America and has a significant impact on developing countries ^2^. Annual DENV infections are estimated at 100–390 million per year ^3^ and dengue is ‘re-emerging’ mostly due to the expansion of the geographical range of the principal mosquito vector, *Aedes (Ae.) aegypti*, through globalization and climate change ^2,4^. ZIKV is a flavivirus related to dengue virus (DENV) and historically thought to be transmitted by *Ae. aegypti*. Local transmission of ZIKV in the Americas was first reported in early 2014 and 22 countries and territories have now been identified to have autochthonous transmission ^5^. YFV is also transmitted by *Ae. aegypti* and can result in large urban outbreaks and rapid spread to distant locations ^6^. Yellow fever is now endemic in Central American countries and in several Caribbean Islands ^7^. CHIKV is an alphavirus transmitted by *Ae. albopictus* (and to a lesser extent by *Ae. aegypti*) and has spread globally with outbreaks in the mid 2000s in the Indian Ocean and India and even in Europe in 2007 ^8^. Transmission of CHIKV has also been seen recently in the Americas and this rapid geographical expansion (in a similar way to DENV) is likely due to the expanding habitat of the mosquito vectors ^4^.

Outbreaks of arboviral diseases including DENV ^9^, YFV ^7^, CHIKV ^10^ and ZIKV ^11^ have recently occurred in the Caribbean. The possibility of additional recent arbovirus transmission in the Caribbean must be considered given some infections result in nearly indistinguishable clinical symptoms. For example, Mayaro virus (MAYV) is an alphavirus closely related to CHIKV and has resulted in sporadic outbreaks in South America ^12^. MAYV transmission is restricted to South and Central America where it is thought that non-human primates act as reservoir hosts and *Haemogogus* mosquitoes (eg. *H. janthinomys*) found in sylvatic jungle environments are responsible for human cases. Although human cases are strongly correlated with exposure to forest environments, urban transmission of MAYV must be considered given the association of cases and major cities infested with *Ae. aegypti* ^13^. As the Caribbean is a destination for many international tourists, surveillance is needed for individual Caribbean islands to determine the risk of facilitating the spread of arboviral diseases. In particular, arboviruses transmitted by *Ae. aegypti* are considered important given that prevention predominantly relies on mosquito vector control. *Ae. aegypti* was first identified in the Caribbean Islands in 1864 ^14–16^ as has remained present despite the Pan American Health Organization (PAHO) mosquito control campaign in the 1940s-1960s that was launched to eliminate urban yellow fever. *Ae. aegypti* was successfully eradicated in many countries including Brazil, Mexico and Guatemala ^17^ but eradication was not achieved in other countries such as the USA, Suriname, Guyana, French Guyana, Venezuela and the Caribbean Islands. As the eradication campaign deteriorated in the early 1970s and 1980s, many countries became re-infested with Ae. *aegypti* ^18,19^ and the geographical expansion of *Ae. aegypti* with urbanization resulted in the introduction of DENV to many countries ^20,21^. With the exception of YFV, there are no currently available treatments or vaccines for arboviral diseases transmitted by *Ae. aegypti* and *Ae. albopictus*. Disease control is currently limited to traditional vector control strategies that rely on insecticides or destruction of larval breeding sites. In most DENV-endemic countries, ultra-low volume space spraying is recommended only during dengue outbreaks. However, widespread insecticide resistance has developed in *Ae. aegypti*, including high pyrethroid resistance rates in South America ^22^ and further north in the Caribbean ^23^.

The volcanic island of Saint Lucia is located midway down the Eastern Caribbean Chain between Martinique and Saint Vincent and north of Barbados (**Figure 1**). The first cases of dengue in Saint Lucia were recorded in the 1980s and following Hurricane Thomas in 2011 another outbreak occurred ^24^. CHIKV was first introduced to Saint Lucia in 2014 ^25,26^ but despite these outbreaks of major mosquito-borne arboviruses, vector surveillance has been limited and the last documented studies were carried out in 1976 ^14–16^. The landscape of Saint Lucia in many areas has changed over the past 40 years due to natural disasters and urbanisation, which has likely changed the distribution of arbovirus vectors. As the density and habitats of Ae. *aegypti* have expanded both in urban and rural areas of many tropical countries, we conducted an initial survey of *Ae. aegypti* and *Ae. albopictus* distribution and analysed any environmental parameters that were associated with the presence of these species. Female mosquitoes were screened for medically important arboviruses and other flaviviruses to investigate whether there was any evidence of infection.

**Figure 1.**
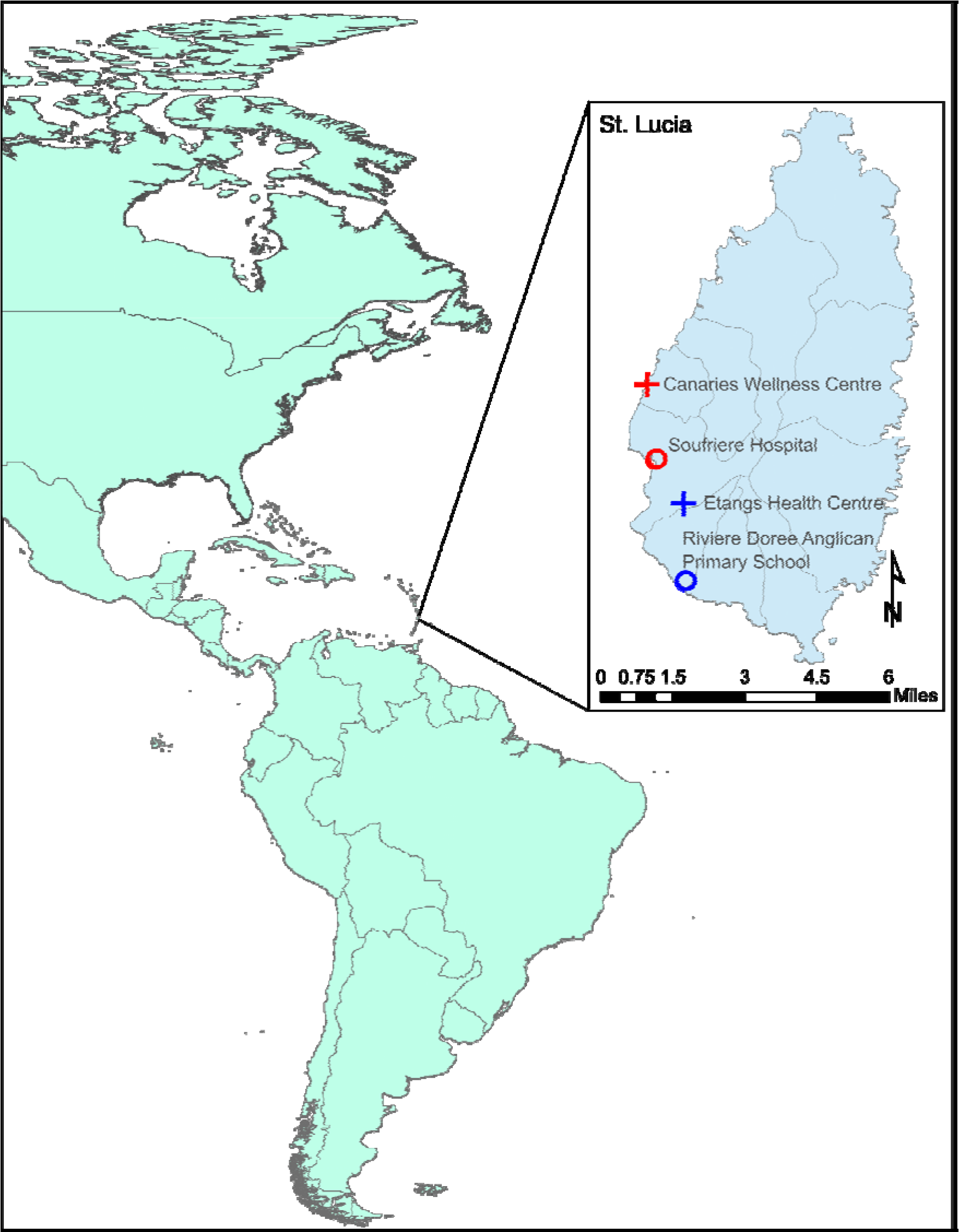
Sampling locations of the longer-term mosquito traps on the island of Saint Lucia used throughout the duration of the study (July 2015). Inset: a representative BG Sentinel 2 trap placed in the Des Cartier rainforest (a temporary trap location).

## METHODS

### Study Sites and mosquito collection

Mosquito collections were carried out on the island of Saint Lucia (Latitude 14.0167°N, Longitude 60.9833°W) in July 2015. Saint Lucia has a population of ~166,000 people and is 27 miles long and 14 miles wide with forest covering 77% of the island. The tropical climate includes a dry season (December to June) and a wet season (July to November). Biogents (BG) Sentinel 2.0 mosquito traps baited with BG lure® were used at various sites across the island (supplementary figure 1) during the beginning of the wet season. Site selection was undertaken based on geographical and environmental variation in urban and semi-urban areas across the island and factors based on island topology including forested areas, brackish water bodies, fresh water bodies and mangrove habitats in communities with previously high mosquito numbers recorded gathered from local knowledge. In some locations traps were placed inside houses. Four permanent traps were connected to power supplies at Canaries Wellness Centre, Soufriere Hospital, Etangs Wellness Centre and the River Doree Anglican Primary School (Figure 1) and traps were run for a period of 22-24 days with mosquitoes collected at 24-hour intervals. Four temporary traps powered by Power King®12 volt batteries were deployed at various locations across the island (supplementary figure 1) to collect mosquitoes over a 24-hour period. Lascar easy log USB data logger 2’s were placed in permanent traps to record humidity and temperature at hourly intervals. Garmin GPS coordinators were used to determine co-ordinates of both permanent and temporary traps. Trapped mosquitoes were collected, killed on ice for morphological identification to identify individuals belonging to the *Aedes* genus. Larval dipping was undertaken at Soufriere Town, Choiseul Village, Marisule and Gros Islet to sample immature stages (larvae/pupae) from domestic containers (e.g. tanks and drums, discarded containers and tires). Immature stages were reared and allowed to emerge in mosquito cages. Individual mosquitoes that were identified by morphology to be *Ae. aegypti* were placed in RNAlater and stored at −20°C to preserve RNA for molecular analysis.

### RNA extraction and PCR analysis of adult *Ae. aegypti*

*Ae. aegypti* adult female mosquitoes were pooled according to trap location and date of collection (1-3 females/pool) and RNA was extracted using Qiagen 96 RNeasy Kits according to manufacturer’s instructions and a Qiagen Tissue Lyser II (Hilden, Germany) with 3mm stainless steel beads to homogenise mosquitoes. RNA was eluted in 45 μl of RNase-free water and stored at −70°C. A Qiagen QuantiTect Kit was first used to remove any genomic DNA co-purified during the RNA extraction protocol and then reverse transcription was performed to generate cDNA from all RNA extracts using manufacturer’s instructions. Confirmation of species identification was undertaken using an Internal transcribed spacer 1 (ITS1) real time PCR assay that discriminates between *Ae. aegypti* and *Ae. albopictus* ^27^. Arbovirus screening included the major arboviruses of public health importance, suspected or having the potential of being transmitted by *Ae. aegypti* / *Ae. albopictus* in the Caribbean: DENV, CHIKV, ZIKV, YFV and MAYV (table 1). In addition, Pan-Flavivirus PCR screening was undertaken that allows simultaneous detection of numerous flaviviruses using a conserved region of the NS5 gene ^28^. PCR reactions for all assays except ZIKV were prepared using 5 µl of Qiagen SYBR Green Master mix, a final concentration of 1 µM of each primer, 1 µl of PCR grade water and 2 µl template cDNA, to a final reaction volume of 10 µl. Prepared reactions were run on a Roche LightCycler® 96 System and PCR cycling conditions are described in table 1. Amplification was followed by a dissociation curve (95°C for 10 seconds, 65°C for 60 seconds and 97°C for 1 second) to ensure the correct target sequence was being amplified. ZIKV screening was undertaken using a Taqman probe based assay ^29^. PCR results were analysed using the LightCycler® 96 software (Roche Diagnostics). Synthetic long oligonucleotide standards of the amplified PCR product were generated in the absence of biological virus cDNA positive controls and each assay included negative (no template) controls.

**Table 1.**
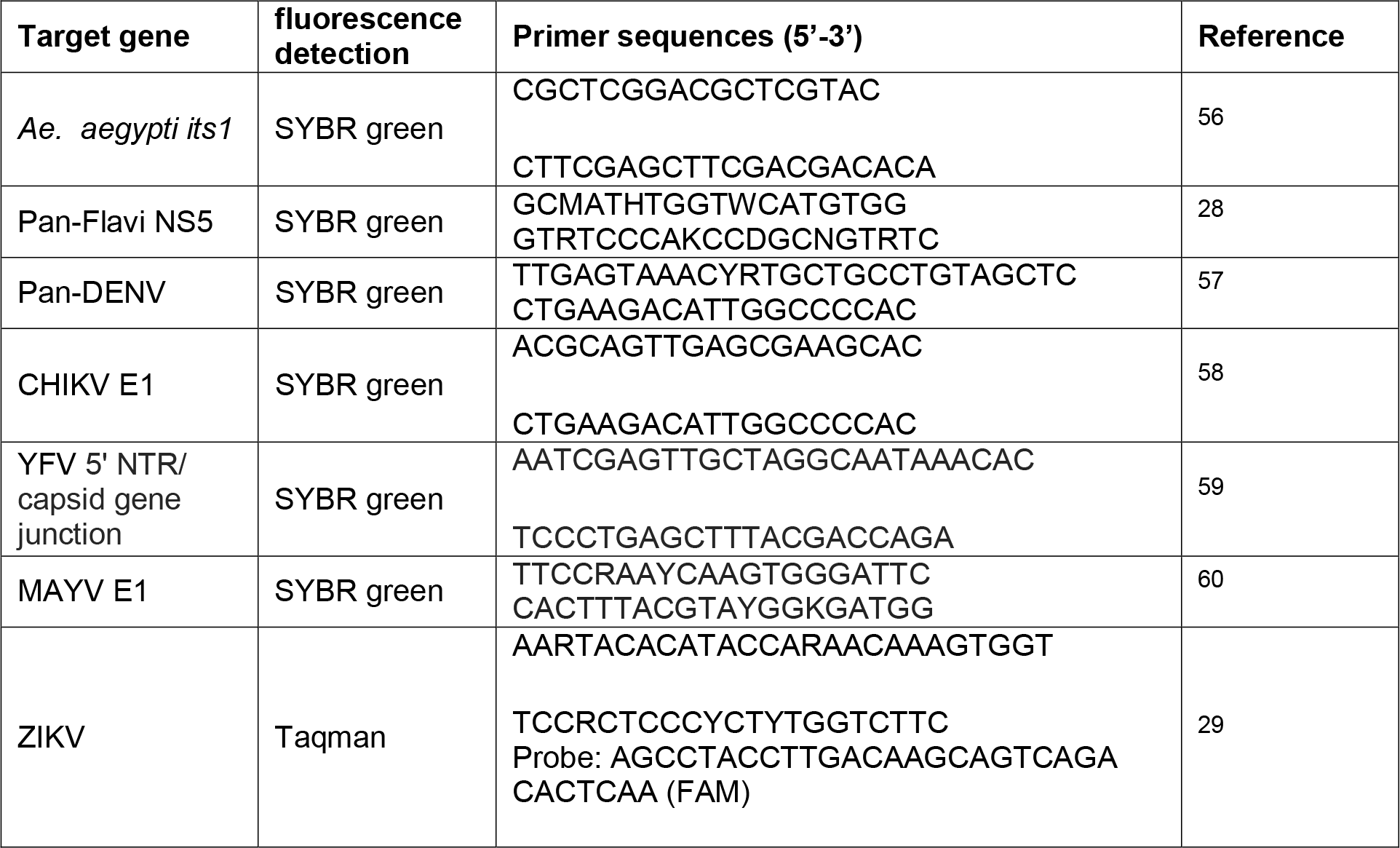
PCR gene targets and primer sequences for the screening analysis undertaken on *Ae. aegypti* mosquito cDNA.

### Sanger sequencing and phylogenetic analysis

Pan-flavi PCR products were submitted to Source BioScience (Source BioScience Plc, Nottingham, UK) for PCR reaction clean-up, followed by Sanger sequencing to generate both forward and reverse reads. Sequencing analysis was carried out in MEGA7 ^30^ as follows. Both chromatograms (forward and reverse traces) from each sample was manually checked, edited, and trimmed as required, followed by alignment by ClustalW and checking to produce consensus sequences. Consensus sequences were used to perform nucleotide BLAST (NCBI) database queries. Maximum Likelihood phylogenetic trees were constructed from Sanger sequences as follows. The evolutionary history was inferred by using the Maximum Likelihood method based on the Tamura-Nei model ^31^. The tree with the highest log likelihood in each case is shown. The percentage of trees in which the associated taxa clustered together is shown next to the branches. Initial tree(s) for the heuristic search were obtained automatically by applying Neighbor-Joining and BioNJ algorithms to a matrix of pairwise distances estimated using the Maximum Composite Likelihood (MCL) approach, and then selecting the topology with superior log likelihood value. The trees are drawn to scale, with branch lengths measured in the number of substitutions per site. Codon positions included were 1st+2nd+3rd+Noncoding. All positions containing gaps and missing data were eliminated. The phylogeny test was by Bootstrap method with 1000 replications. Evolutionary analyses were conducted in MEGA7 ^30^.

### Statistical analysis

Count data analysis was conducted using a generalized linear model because the response variable (mosquito counts) had a non-normal error distribution. Models were run using Stata MP (version 14, Stata Corp, College Station, TX, USA). Both Poisson and negative binomial link functions were used in analysis, with the superior model identified from visual inspection of fits (**Supplementary figure 2**). A univariate analysis included elevation, humidity and temperature as continuous explanatory variables, and urbanisation level (urban or semi-urban) as a factor. Incident Rate Ratios (and corresponding 95% confidence intervals) were calculated.

## Results

A total of 3,701 adult mosquitoes were collected across the island of Saint Lucia over a four-week period using BG sentinel 2 traps (**Table 2**). *Culex* was the dominant genus, comprising 78.7% of the total mosquitoes collected and the remaining 21.3% were morphologically identified as species within the *Aedes* genus. No *Ae. albopictus* females were collected in any of the locations despite traps being placed in diverse ecological settings. *Ae. aegypti* adults were collected in 26/46 trap locations with the largest number of females being collected at Soufriere Hospital (n=196) and Canaries Wellness Centre (n=93) where permanent traps were running for the duration of the collection period. The average number of female *Ae. aegypti* collected over a 24-hour period across all trap locations was 3.09. A particularly high number of *Ae. aegypti* were collected during a 24-hour period from Dugard (47 females and 10 males comprising 47.5% of the total collection) (**Figure 2**, **supplementary figure 1**) using a trap placed indoors in a semi-urban area. In contrast, low numbers of *Ae. aegypti* were collected using the permanent trap at River Dorree Anglican Primary School with *Ae. aegypti* compromising 0.5% (n=5) of the collection and an average of 0.13 female mosquitoes per 24 hours of trapping.

**Table 2.**
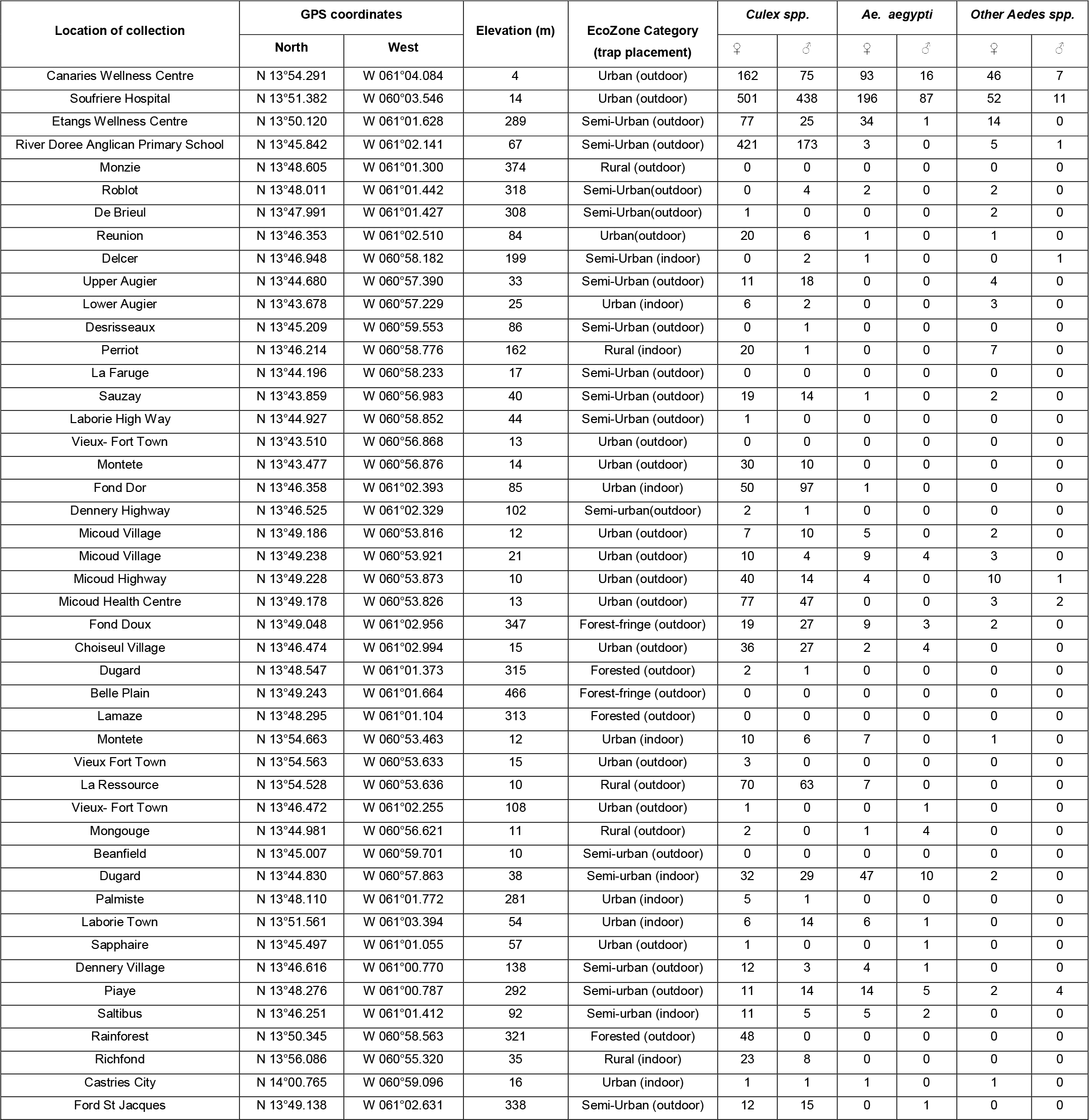
Collection site locations and characteristics with total numbers of adult mosquitoes collected from each site.

**Figure 2.**
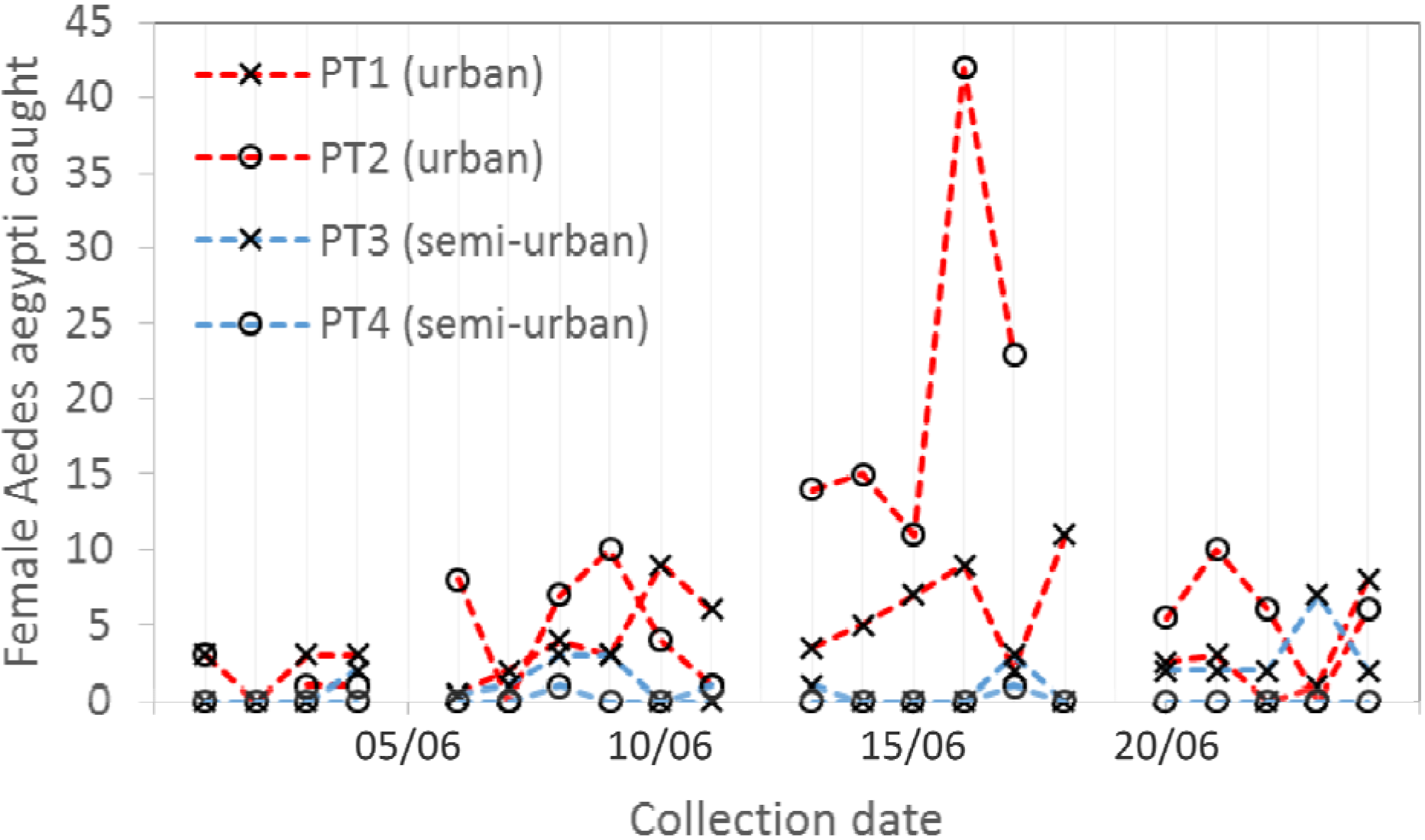
Population dynamics of local female *Aedes aegypti* mosquitoes caught from the four longer-term traps positioned in field sites detailed in Figure 1.

A generalized linear model (GLM) was used to analyse the associations between the counts of female *Ae. aegypti* (combining counts from both the temporary traps and permanent traps) and four independent variables: peak daily temperature, peak daily humidity, trap elevation and ecological zone (semi-urban or urban). Plotting count frequencies against alternative, competing models assuming either a Poisson or a negative binomial distribution clearly demonstrated the superiority of a negative binomial model in fitting the data distribution (**supplementary figure 2**). Exponentiation of the coefficients resulting from a GLM (negative binomial family) produced the incidence rate ratio (IRR) associated with the independent variables. Here, IRR can be interpreted as the ratio of counts per trap-day associated with the tested variable. These are described along with 95% confidence intervals in **Table 2**. No significant association was found with temperature. Because previous studies have shown a non-monotonic association between *Ae. aegypti* and temperature (i.e. *Ae. aegypti* thrive at a non-trivial optimal temperature) ^32^ we subsequently attempted to fit a more complex (quadratic) function between these variables but this did not improve model fit (data not shown). Higher counts were significantly associated with higher elevation although the effect size was small; and semi-urban settings yielded female mosquito counts per trap-day that were 5-fold lower than urban settings. We tested for interactions between all covariates but none were found to be significant.

A sub-sample of adult female *Ae. aegypti* mosquitoes collected from BG traps were screened for arboviruses (**Table 3**). No evidence was seen for infection of the major medically important arboviruses that have historically been transmitted by *Ae. aegypti*. However, the presence of a novel flavivirus closely related to cell fusing agent virus (CFAV) was detected (**Figure 3**) in 17.8% (8/45) individuals screened from Soufriere Hospital, 20% (3/15) of individuals screened from Etangs Wellness Centre, 33% (1/3) screened from Micoud Village, 50% (1/2) individuals from Micoud Highway and 50% (2/4) individuals from Piaye. We also detected this novel flavivirus in adult females that have been reared from larval collection (n=10). Phylogenetic analysis reveals this flavivirus is an insect specific flavivirus (ISF) clustering with other ISFs and separate from pathogenic flavivirus such as DENV, ZIKV and YFV (**Figure 4**).

**Table 3.**
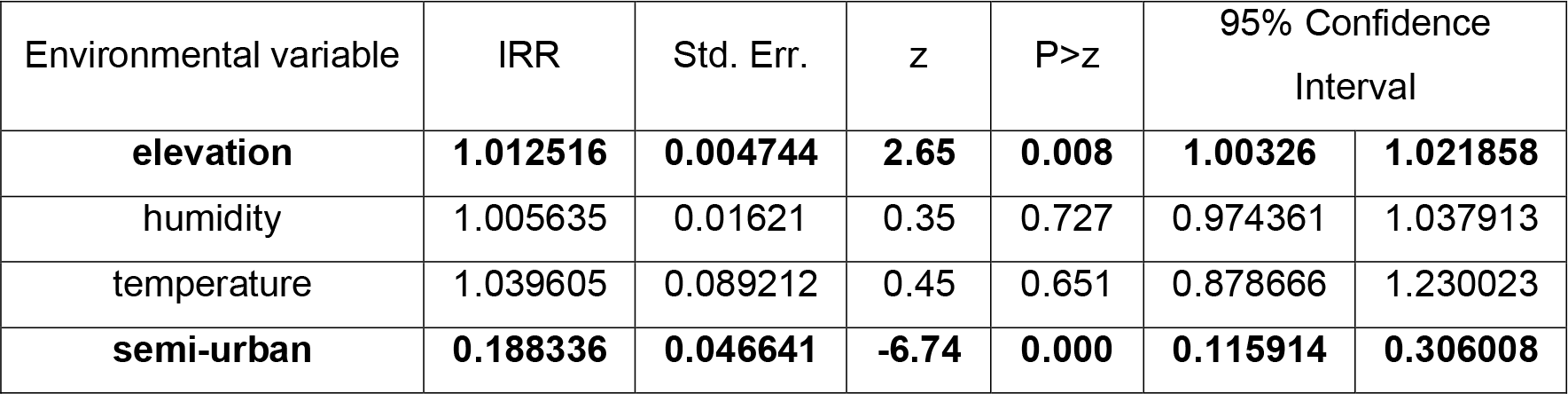
Incidence Rate Ratios (IRR) and corresponding 95% confidence intervals resulting from univariate generalized linear models with negative binomial link function.

**Figure 3.**
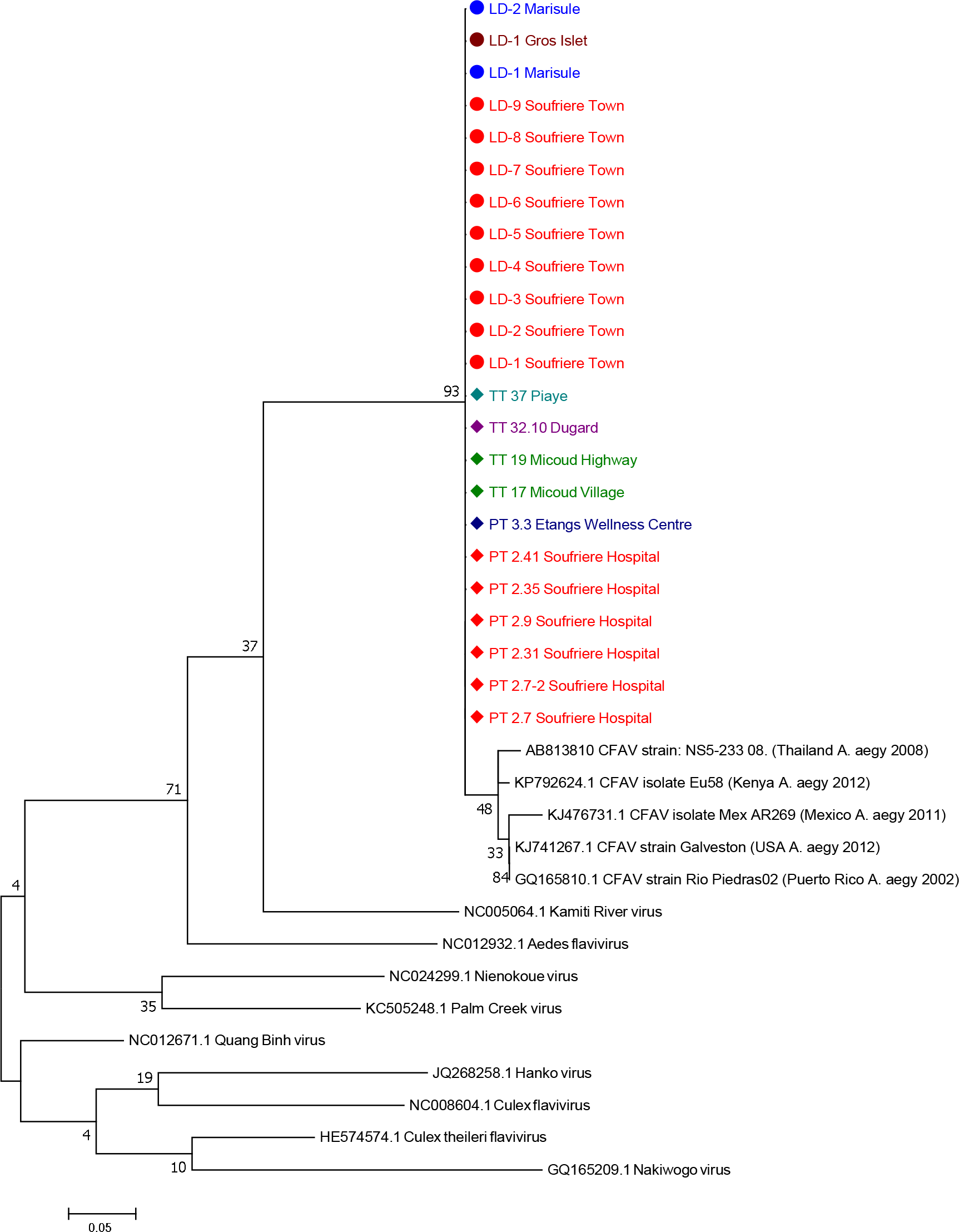
Maximum Likelihood molecular phylogenetic analysis of Pan-flavi NS5 sequences from field-collected *Ae. aegypti* mosquitoes. The tree with the highest log likelihood (−1077.12) is shown. The tree is drawn to scale, with branch lengths measured in the number of substitutions per site. The analysis involved 37 nucleotide sequences. There were a total of 124 positions in the final dataset.

**Figure 4.**
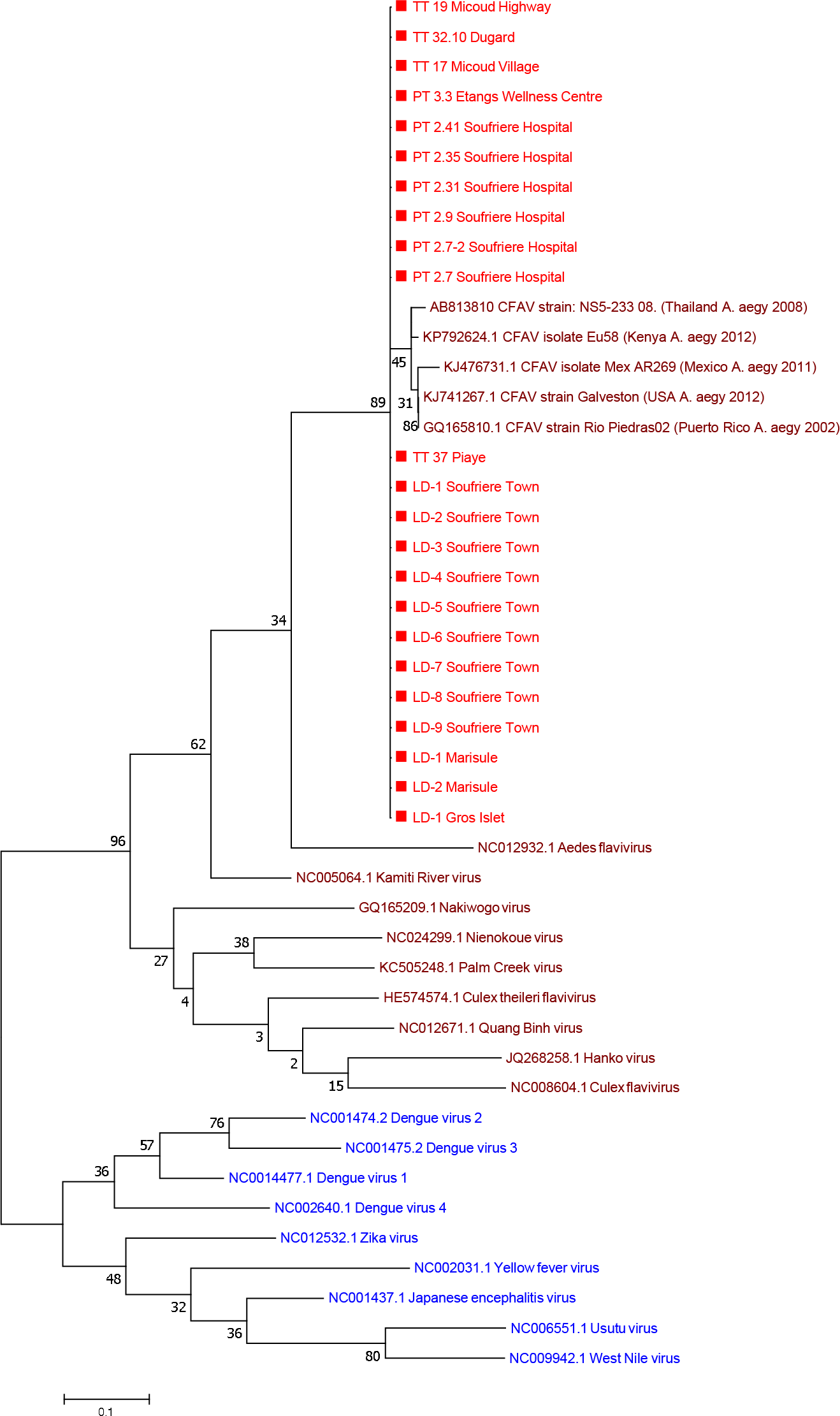
Maximum Likelihood molecular phylogenetic analysis of Pan-flavi NS5 sequences showing the novel flavivirus clustering with insect specific flaviviruses. The tree with the highest log likelihood (−1951.89) is shown. The tree is drawn to scale, with branch lengths measured in the number of substitutions per site. The analysis involved 46 nucleotide sequences. There were a total of 124 positions in the final dataset.

## Discussion

Entomological indices including the abundance of adult mosquitoes are often used to assess the risk of disease transmission and this, in turn, influences vector control strategies. The lack of surveillance studies, to our knowledge, for major vectors of arboviruses of public health importance on the island of Saint Lucia needed to be addressed, given the recent outbreaks of arboviruses such as DENV, CHIKV and ZIKV in the Caribbean and surrounding regions. The principle vector of these arboviral diseases, *Ae. aegypti*, is highly invasive and is now present in much of the Americas including the US ^5^. In this study, we collected adult mosquitoes to determine the geographical distribution of *Ae. aegypti* across the island of Saint Lucia and to determine any correlation with environmental variables. BG sentinel 2 traps were selected as these traps were redesigned to provide increased durability in field conditions and were recently shown to be effective at trapping *Aedes* species ^33^. The durability of traps was particularly important for the four permanent traps that were used for approximately 24 days. Adult collections indicate that *Ae. aegypti* is present throughout the Island of Saint Lucia and population densities are significantly higher in urban areas compared to semi-urban or rural settings. We also demonstrated that adult counts were positively correlated to elevation.

The trapping of a high number of *Ae. aegypti (*47 females and 10 males) during a 24-hour period from Dugard using a trap placed indoors in a semi-urban area suggests an interesting behavioural observation. The biology and behaviour of *Ae. aegypti* in the Caribbean has not been extensively studied but previous work on a Trinidad strain using human landing catches revealed the periodicity consisted of 90% arriving during daylight and twilight and 10% during the night ^34^. This study included both urban and rural sites and a consistently larger number of mosquitoes were collected outside vs. inside houses. Light intensity was also significantly correlated with mosquito landing patterns ^34^. The trapping of *Ae. aegypti* inside houses in Saint Lucia could indicate a change in behavior with mosquitoes biting indoors during the night in houses with lights on (an anecdotal observation that occurred during our study). Indoor resting of *Ae. aegypti* has recently been documented in Mexico ^35^ which has implications for control methods. Other studies in the Caribbean have also shown that high temperatures in open environments can result in *Ae. aegypti* breeding in underground sites ^36^ and indoor oviposition has been demonstrated ^37,38^.

Confirmation of the presence of *Ae. aegypti* on the island of Saint Lucia is not particularly surprising given this species is widespread throughout the Caribbean and is now widespread in the Americas ^4,5^. The association with urban environments in Caribbean Islands is seen with the most common breeding sites being drums/barrels, uncovered tanks and cisterns, brick holes, flower pots, used tyres and utility manholes ^39,40^. Saint Lucia now provides the ideal environment for *Ae. aegypti* due to recent changes in the climate. The El Nino period in 2009 - 2010 introduced dry hot periods and provided an environment that was not conducive for mosquito production ^40^. Water conservation has become a critical issue for Saint Lucia and the majority of the water supply comes from surface runoff collected in rivers, streams and dams. Rain water is collected and stored haphazardly and inappropriately in various containers such as water tanks, drums, and buckets, creating ideal breeding grounds for this species ^1,40^. This study was undertaken during the commencement of the wet season with the average rainfall in June and July 2015 being 37.1mm and 175.8mm respectively. This indicates that greater mosquito abundance is likely throughout later stages of the wet season and follow-up studies should be undertaken to determine this. Clearly climatic patterns resulting in unpredictable rainfall will provide ideal breeding grounds (unpolluted water in artificial and natural containers) for *Ae. aegypti* in Saint Lucia ^36^.

*Ae. albopictus* was not identified in the mosquitoes collected in our study despite many traps being set in or near forested areas. Although the range of *Ae. albopictus* has expanded to Europe, USA and many South American countries ^4^, this species has only been found in the Eastern Caribbean ^40^. However, recent outbreaks of CHIKV on Saint Lucia and neighbouring Caribbean islands suggest that there might be a possibility that *Ae. albopictus* may also play a role in the spread of the disease. Although not confirmed in most Caribbean islands, the Dominican Republic was the first country to confirm the presence of Ae. *albopictus* ^41^. With *Ae. albopictus* present in the US to the north and the Cayman Islands to the south, Saint Lucia is clearly considered at risk for establishment of *Ae. albopictus* ^42^. *Ae. albopictus* has also been shown to harbour Eastern equine encephalitis virus (EEEV) ^43^, highlighting the potential transmission risk of additional arboviruses. The traditional ways of importing *Ae. albopictus* through the trade of tyres is also a possible source of introduction for this species. Although Saint Lucia has signed onto the International Health Regulations (IHR) 2005, to prevent and control the international spread of disease, port surveillance systems are not fully implemented and might not be sufficient to monitor containers present on ships and ensure that they are fumigated before they arrive in port. Saint Lucia is also faced with the problem of tyre disposal where there is no functional shredding equipment, which is of great concern, particularly so because tyres in landfills are in close proximity to urban communities.

The detection of a novel flavivirus closely related to CFAV in diverse ecological populations of *Ae. aegypti* across the island of Saint Lucia suggests the potential for undiscovered viruses in the Caribbean. A large study was undertaken in Trinidad screening more than 185,000 mosquitoes representing 46 species and 85 different viruses were isolated ^44^. The isolation of Mucambo virus (MUCV), a Venezuelan Equine Encephalitis complex subtype IIIA), follows a history of isolating alphaviruses from mosquitoes in Trinidad^45^. A potentially novel strain of CFAV was discovered in *Ae. aegypti* populations from Mexico ^46^ and CFAV was detected in *Ae. aegypti* populations from Kenya ^47^. Interestingly, CFAV infection significantly enhanced replication of DENV (and vice versa) in *Ae. aegypti* Aag2 mosquito cells ^48^. CFAV has been shown to be vertically transmitted in *Ae. aegypti* lab colonies suggesting the possibility of using CFAVs and closely related ISFs for control of medically important arboviruses ^49^. The presence of insect-specific viruses in *Ae. aegypti* might be underestimated given a recent study suggested up to 27 insect-specific viruses (23 currently uncharacterized) in populations from Cairns (Australia) and Bangkok (Thailand) ^50^. The question remains as to whether insect specific viruses like CFAV have not yet gained the ability to infect vertebrates and therefore become arboviruses or whether they have lost this ability ^51^. Phylogenetic studies focussed on the E gene of flaviviruses would suggest CFAV is a basal lineage that diverged prior to the separation of mosquito and tick-borne flaviviruses ^52^. Our results indicate the presence of a flavivirus but it has been shown that some flavivirus genome integrated sequences can be transcribed and therefore cautious must be taken to assume the presence of an active flavivirus infection ^53^. The impact of arboviral diseases is increasing due to the expanding geographical range of many mosquito species, particularly *Ae. aegypti* and *Ae. albopictus*. As most arboviral diseases occur in sporadic epidemics, vector control options are often limited to the use of insecticides that are becoming less effective due to insecticide resistance. As re-emerging arboviral diseases such as DENV and ZIKV continue to spread geographically, the fight to eradicate or reduce the transmission potential of *Ae. aegypti* is increasing in importance. Outbreaks of arboviral diseases, including DENV, CHIKV and ZIKV, have a history of occurring in small tropical islands. ZIKV emerged for the first time outside of Africa and Asia in Yap State in Micronesia and then a large outbreak in French Polynesia was followed by transmission in other Pacific islands ^54^. Small Islands Developing States and territories (SIDS) such as Saint Lucia are particularly vulnerable to arboviral disease outbreaks for several reasons ^55^. Natural disasters are more frequent and these change the geographical landscape allowing rapid mosquito proliferation. SIDS often lack safe water supplies and sanitation and local governments have limited resources to undertake vector control and manage outbreaks. An increasing ability for travel between SIDS and continental regions facilitate the spread of arboviruses to previously unexposed populations. For these reasons, surveillance strategies need to be monitored, risk areas need to be mapped out and epidemic trends recorded for predicting future outbreaks. For the Caribbean island of Saint Lucia, further research is needed to determine the diversity of current mosquito species and this should be extended to the neighbouring smaller Caribbean islands.

## Acknowledgements

We would like to thank all the villagers who participated in the fieldwork. Funding was provided by the London School of Hygiene and Tropical Medicine, Bayer Pharmaceuticals and a Wellcome Trust /Royal Society grant awarded to TW (101285/Z/13/Z): http://www.wellcome.ac.uk; https://royalsociety.org. The funders had no role in study design, data collection and analysis, decision to publish, or preparation of the manuscript.

## Author contributions statement

All authors contributed to the design of the study. MW and LW performed mosquito collections. CLJ, MW, LW and TW performed molecular analysis of samples. CLJ performed Sanger sequencing and phylogenetic analysis. CLJ, MW, LY and TW undertook data analysis. All authors read and approved final version of the manuscript.

## Availability of data and material

All data generated or analysed during this study are included in this article.

## Supplementary information

**Supplementary Figure 1:**
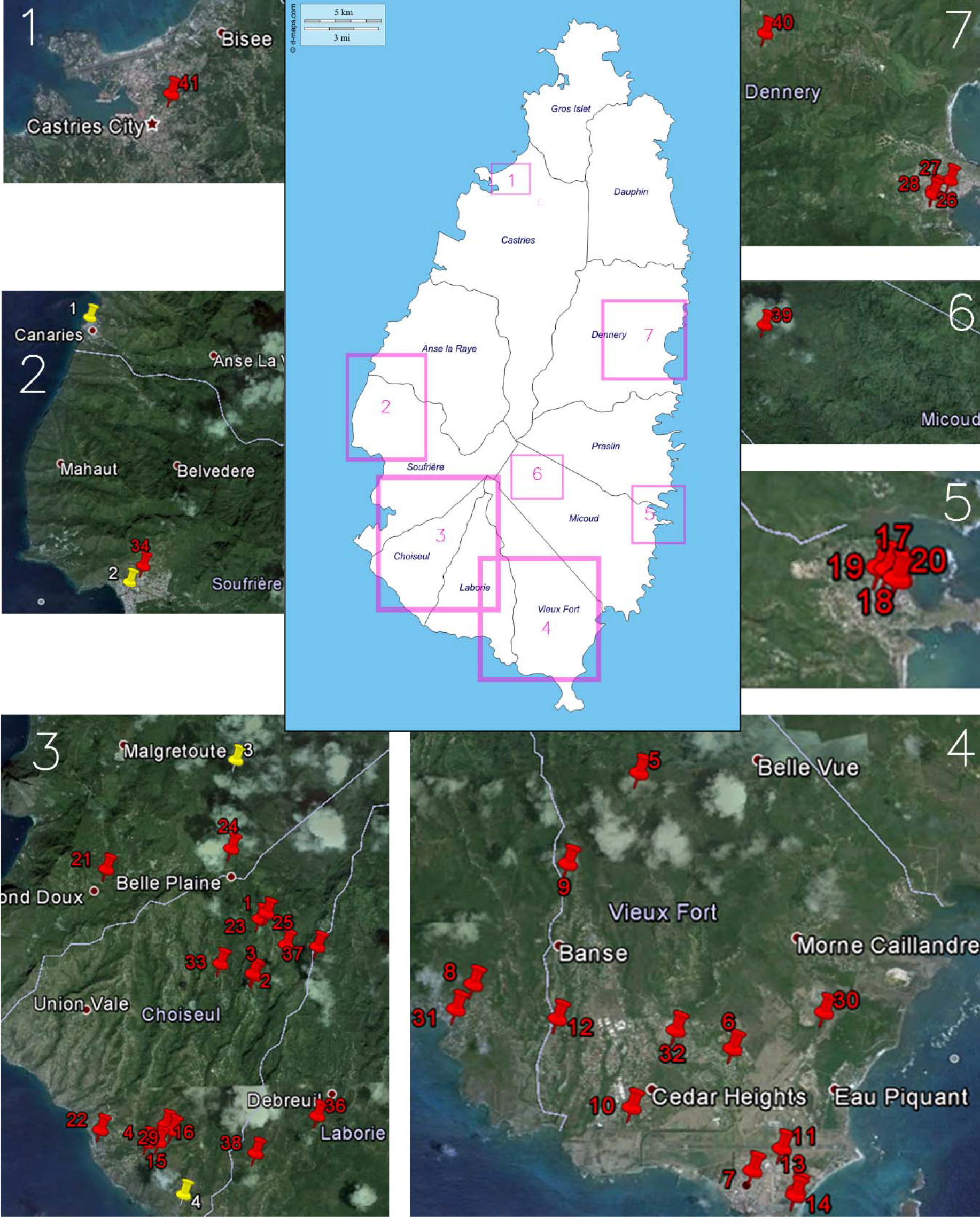
Map of Saint Lucia showing the location of BG Sentinel 2 traps used in the study. GPS coordinates were plotted using Google Earth. Red pins represent locations of temporary traps and yellow pins permanent traps.

**Supplementary Figure 2.**
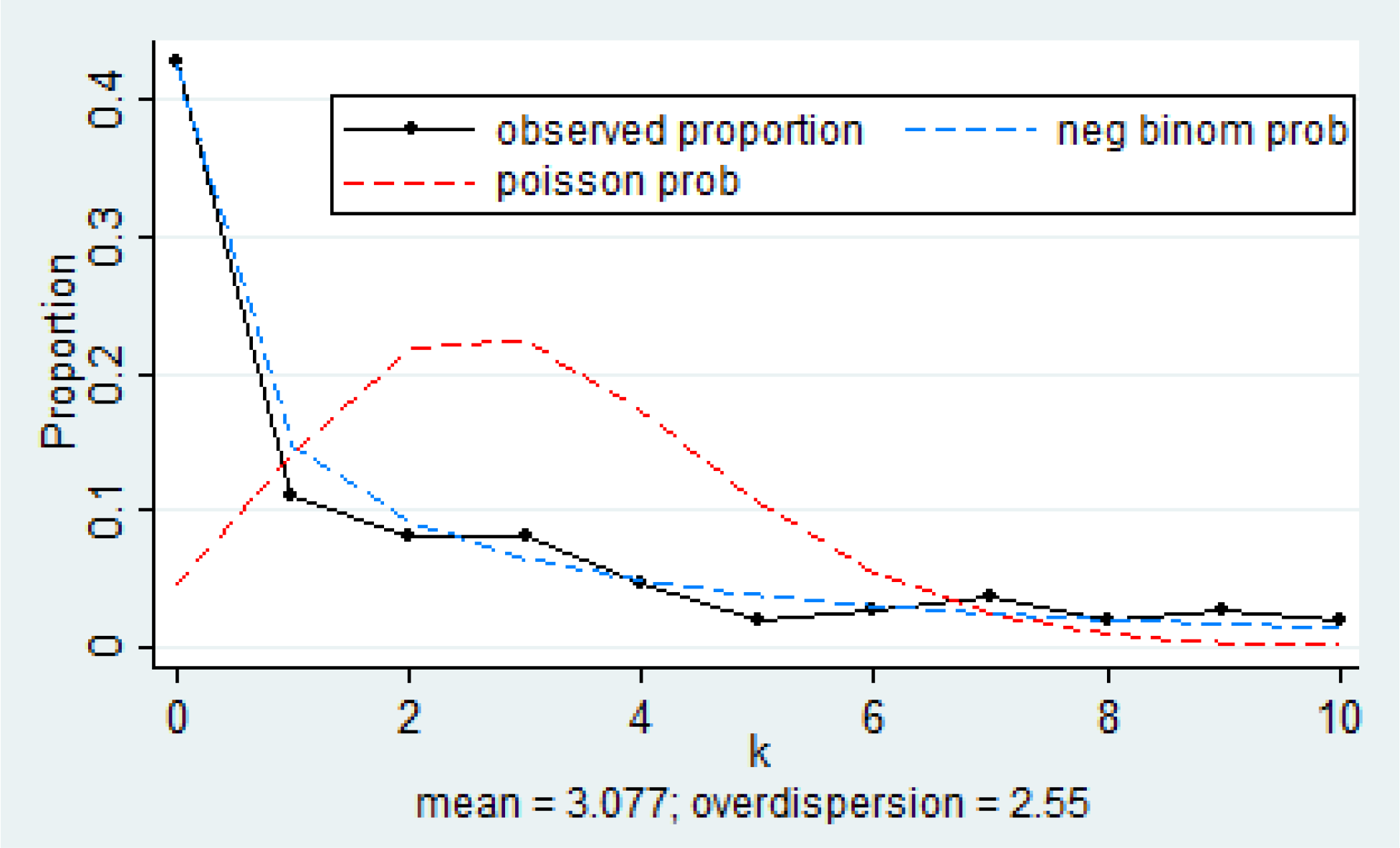
The observed proportions along with the Poisson and negative binomial probabilities for the count type variable (using ‘nbvargr’ function in Stata). The Poisson probabilities are computed using an estimate of the Poisson mean. The negative binomial probabilities use the same mean and an estimate of the over dispersion parameter.

